# High genomic and phenotypic diversity among *Candida dubliniensis* isolates from the oral cavity of persons living with HIV

**DOI:** 10.1101/2025.10.23.684142

**Authors:** Verónica A. Dubois, Marina Marcet-Houben, Diego Fuentes-Palacios, Ewa Ksiezopolska, Laura A. Gliosca, Toni Gabaldón

## Abstract

The opportunistic yeast pathogen *Candida dubliniensis* is frequently misidentified as its closest relative *Candida albicans*, due to their high phenotypic similarity. Although *Candida dubliniensis* is an important cause of fungal infections, especially in the oral cavities of immunocompromised patients, its phenotypic and genomic variability remains poorly understood. We addressed this knowledge gap by performing a comprehensive phenotypic and genomic characterization of 33 *C. dubliniensis* oral isolates from people living with HIV in Argentina. Our results uncovered significant phenotypic heterogeneity—observed both inter- and intra-patient—impacting clinically relevant traits such as drug susceptibility, stress sensitivity, filamentation, and hypoxic growth. Our isolates expanded the previously known MLST-based genetic diversity of the species. Leveraging whole-genome sequencing, we resolved intricate evolutionary relationships, proposed a more robust MLST scheme for future typing, and identified potential antifungal resistance-causing variants. Collectively, this work represents the most comprehensive population genomics characterization of *C. dubliniensis* to date, establishing a foundation for improved surveillance and diagnostic strategies to fully define its role in both oral and systemic infections.

**Author Summary:** In this study we uncover hidden genetic and morphological diversity in *Candida dubliniensis,* a poorly known opportunistic yeast pathogen related to *Candida albicans*, by characterising a collection of oral isolates from persons living with HIV in Argentina. Our findings provide the most detailed genetic and phenotypic snapshot of this pathogen to date, providing a much-needed foundation to develop better diagnostic tools and effective surveillance strategies to address infections caused by this overlooked threat.

## INTRODUCTION

*Candida dubliniensis* is an opportunistic yeast pathogen that is the closest relative of *Candida albicans* [1, 2]. Traditionally, the differentiation between these two species relied on the assessment of morphological and metabolic differences [3]. However, it is currently well known that diagnosis based on such phenotypes can lead to underdiagnosis of *C. dubliniensis*, since only molecular methods allow a precise differentiation from the more prevalent *C. albicans* [4, 5]. Numerous studies have documented a global distribution of *C. dubliniensis,* and existing surveys that consider this species generally report a relatively lower prevalence with respect to other *Candida* species, ranging between 0.2% and 6% [3]. However, to date there is no accurate global estimate of *C. dubliniensis* prevalence, due to frequent misidentification as *C. albicans* [3, 5–9]. Nevertheless, recent studies have documented an increase in the number of cases of systemic candidiasis caused by *C. dubliniensis* [10–20]. This increase might be related to changes in diagnostic procedures [21–23], or alternatively, to causes similar to those underlying the growing isolation of other non-*Candida albicans* species [24–26].

*C. dubliniensis*, like *C. albicans*, belongs to the CTG clade [8, 27, 28]. Species within this clade express multiple virulence factors that play essential roles at various stages of infection. *C. dubliniensis* exhibits phenotypic plasticity similar to that of *C. albicans* [29–31]. This ability to adapt and grow in different morphologies is considered the most important virulence factor of the clade, as it is crucial for the fungus to quickly adjust to the host’s changing environment. This capability enhances the efficiency of this yeast in both infection and tissue invasion [30, 31]. Many pathogenic species of the CTG clade such as *C. albicans*, *C. dubliniensis*, *Candida tropicalis*, and species from the *Candida parapsilosis* complex primarily reproduce asexually. However, an atypical reproductive cycle known as parasexuality has been identified in *C. albicans* [32]. This process begins with the fusion of cells of opposite mating types, leading to the formation of a tetraploid cell. The subsequent return to a diploid state happens in the absence of meiosis, via a process of concerted chromosome loss that stochastically removes one of the two homologous chromosomes of each pair. This mechanism results in genetically diverse progeny and is common in other species without a known sexual cycle [33].

Human colonization by *C. dubliniensis* remains relatively unexplored, particularly when compared to the accumulated knowledge on *C. albicans* [34, 35]. In the immunocompetent population, colonization by *Candida* spp. in the oral cavity ranges from 17% to 75% worldwide [36, 37]. However, it remains uncertain whether the oral cavity, or any specific niche within it, represents the main natural habitat of commensal yeasts [3, 8]. The most crucial factor enabling colonization of a particular niche is adaptation to environmental stress conditions associated with it [38]. Yeasts are capable of altering their metabolism to adapt to environmental conditions and of mounting a combined response to thermal, osmotic, oxidative, nitrosative stress, phagocytosis, and antifungal drugs [39]. However, we still have a very poor understanding about what phenotypic traits enable colonization of humans by *C. dubliniensis*.

In a recent study we discovered some *C. dubliniensis* isolates with morphological characteristics atypical for this species [22], suggesting higher levels of phenotypical diversity than previously anticipated. Moreover, given the scarcity of genetic studies focusing on this species, we have only a very incomplete knowledge on its intrinsic genomic variability. Genome variation is crucial for fungal adaptation, providing the necessary flexibility to adapt to various environments and can ultimately affect virulence-related phenotypes [40, 41]. Given the limited knowledge about this species and its underdiagnosis, we here set out to investigate the genomic and phenotypic variability of oral isolates of *C. dubliniensis*, focusing on their ability to sustain stress conditions that they might encounter in the oral cavity.

## RESULTS

### High phenotypic diversity in stress and drug sensitivity across *C. dubliniensis* isolates

To assess phenotypic diversity in *C. dubliniensis*, we analyzed growth under a panel of stress conditions on a previously available clinical isolate collection (see Materials and Methods, and [22]). This collection comprised 33 isolates from the oral cavity of twelve persons living with HIV, of which eight had confirmed dysbiosis of the periodontal niche—a setting where fungal colonization remains largely underexplored. Considering the clinical relevance of this environment, we sought to investigate growth phenotypes under stress conditions that simulate those encountered in their natural habitat.

We first measured growth in liquid media in control conditions (YPD) or supplemented with various stressors, and obtained different quantitative growth parameters (see Materials and Methods, **Supporting information Fig. SF1 to SF13**). For each condition, including control, our isolates displayed a broad range of growth capacities, as measured by the area under the growth curve (AUC, **Fig. 1A**). Growth under glycerol, sorbitol, SDS or mild osmotic stress (NaCl 0.5M) displayed the broadest range of growth capacities but with average growth similar to that of YPD. In contrast, stressors such as NaCl 0.75M, Dithiothreitol (DTT), Ethanol, Calcofluor White (CFW), Congo Red (CR) or oxidative stress (H₂O₂), and particularly the latter two, caused clear growth defects in most strains. Given the broad growth disparities observed between strains even in control conditions (Fig. 1A), we normalized the growth of each isolate in each stress, as the area under the growth curve (AUC), by its growth in control conditions (YPD), to compute the relative AUC (rAUC) of each isolate, which represents the proportional impact on growth of that particular stress, independently of basal growth differences across strains (see Materials and Methods, **Fig. 1B**). These results confirmed the above-mentioned trends but provided higher resolution in defining stress-specific sensitivities at the isolate level. Importantly, when grouping isolates originated from the same patient, it became apparent that there is a large intra-patient phenotypic heterogeneity. For instance patients 20-0-C, 23-0-C and 24-0-C each harbor strains with highly contrasting phenotypes in terms of sensitivity to oxidative stress. Such intra-patient variability suggests the coexistence of multiple fungal subpopulations within the same host.

**Fig. 1:**
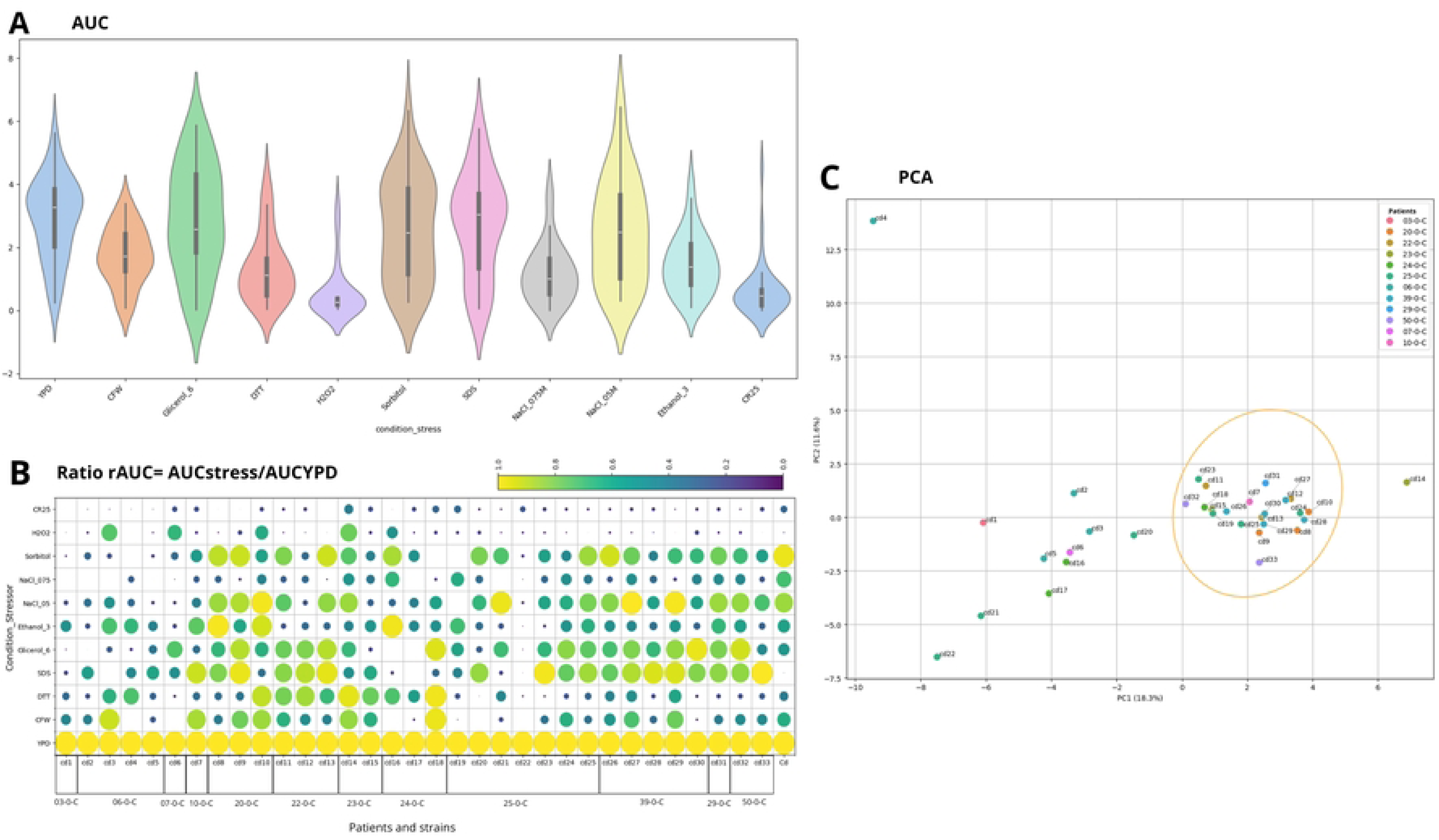
Differential stress responses among clinical isolates assessed by AUC-based metrics and principal component analysis. A) Violin plots showing the distribution of area under the curve (AUC) values across different stress conditions. Each violin represents the distribution of AUC values for all clinical isolates exposed to a specific stressor. The width of each violin reflects the density of strains at a given AUC value. YPD medium was the baseline reference for growth. A marked reduction in AUC (indicating decreased growth) is observed under exposure to hydrogen peroxide (H₂O₂) and Congo Red (CR25), followed by ethanol 3% and NaCl 0.75M. In contrast, growth in the presence of sorbitol 1M, CFW, and glycerol 3% is more similar to YPD, suggesting better tolerance under those conditions. B) Heatmap-style bubble plot showing relative AUC (rAUC) values for each strain under each stress condition, normalized to growth in YPD. Rows correspond to stress conditions and columns to individual clinical isolates, grouped by patient of origin (x-axis). The size and color of each circle represent the magnitude of the rAUC value (see color scale). Larger, yellow circles indicate higher relative tolerance (rAUC ≈ 1), whereas smaller, darker circles indicate increased sensitivity. The plot reveals substantial inter-strain and inter-patient variability in stress resistance profiles. C) Principal component analysis (PCA) of normalized growth parameters across all stress conditions. Each point represents an individual strain, color-coded by patient of origin. PCA was performed using parameters extracted with the G*rowthcurver* package: k, n0, r, t_mid, t_gen, and auc_e. The first two principal components (PC1 and PC2) explain 18.3% and 11.5% of the total variance, respectively. While most isolates cluster closely, indicating overall phenotypic similarity, strains Cd-4-22-21-1-14 are clearly separated in the ordination space, suggesting divergent or unique stress response phenotypes.

To better visualize phenotypic differences, we performed a Principal Component Analysis (PCA), collectively considering six different growth parameters for each of the tested stressors (78 features in total, see Materials and Methods, **Fig 1C**). The two principal axes of the PCA collectively explained 29.9% of the variability. The first axis (18.3%) separated strains across a gradient from higher (left) to lower (right) number of conditions with growth defects, while the second axis (11.6%) captured specific patterns of growth/stress relationships. Most of the isolates clustered in a defined subregion of the PCA (marked with a yellow ellipse), which comprised isolates that have common sensitivities to stresses such as H₂O₂ and NaCl 0.75M, and all have a relatively good growth capacity in NaCl 0.5M, sorbitol, glycerol, and SDS. While some isolates from the same patients are relatively close in this multidimensional analysis, others are scattered across a large area. Interestingly, these include isolates that were obtained not only from the same patient but also from the same periodontal site and at the same date (e.g. isolates from patients 06-0-C, 24-0-C, 25-0-C), highlighting the coexistence of strains with markedly heterogeneous phenotypes.

We next tested these and other stress conditions on solid media using spot assays, which provided consistent results for the shared stresses and confirmed a high level of phenotypic heterogeneity over a larger set of conditions (**Fig. 2**). Solid assays were additionally used to evaluate filamentation by growing strains on spider agar at 30°C and 37°C. Increased filamentation was observed at 37°C, underscoring the role of temperature in morphogenetic responses (**Fig. 3**). Notably, the type strain (CBS7987) failed to filament under any tested condition. Growth in anaerobiosis, which we measured only in solid media, was markedly reduced with respect to aerobiosis, with some isolates exhibiting filamentous morphologies on anaerobiosis depending on the culture medium (16 on YPD, 3 on ASABRU, see **Supporting information Fig. SF14 and SF15**).

**Fig. 2:**
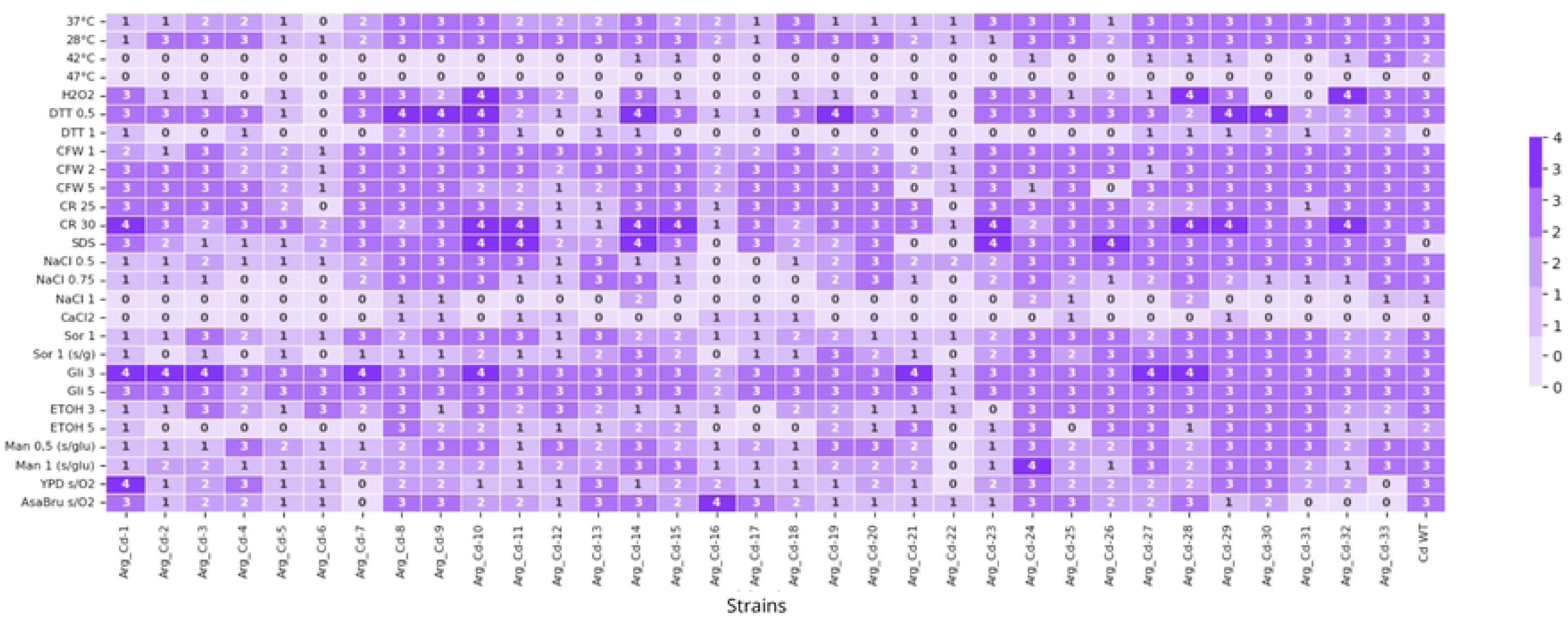
Semi-quantitative categorization of growth in spots of isolates under different stress conditions tested. The number and intensity of the background color indicates the degree of growth, assigning numeric values from 0 to 4 to establish the observed growth level [68]: 0 = no growth (light purple), 1 = minimal growth, 2 = moderate growth, 3 = normal growth (similar to WT strain), 4 = overgrowth (darkest purple). H_2_O_2_ (Hydrogen peroxide), DTT (Dithiothreitol 0.5 and 1 M), CFW (Calcofluor White 1, 2, and 3 g/L), CR (Congo Red 25 and 30 μg/ml), SDS (Sodium dodecyl sulfate), NaCl (Sodium chloride 0.5, 0.75, and 1 M), CaCl_2_ (Calcium chloride), Sor (Sorbitol 1 M), s/g (Without glucose), Gli (Glycerol 3 and 5%), ETOH (Absolute ethanol 3 and 5%), Man (Mannitol 0.5 and 1 M), YPD (Yeast Peptone Dextrose), s/O2 (Without oxygen), ASABRU (Brucella blood agar with hemin).

**Fig. 3:**
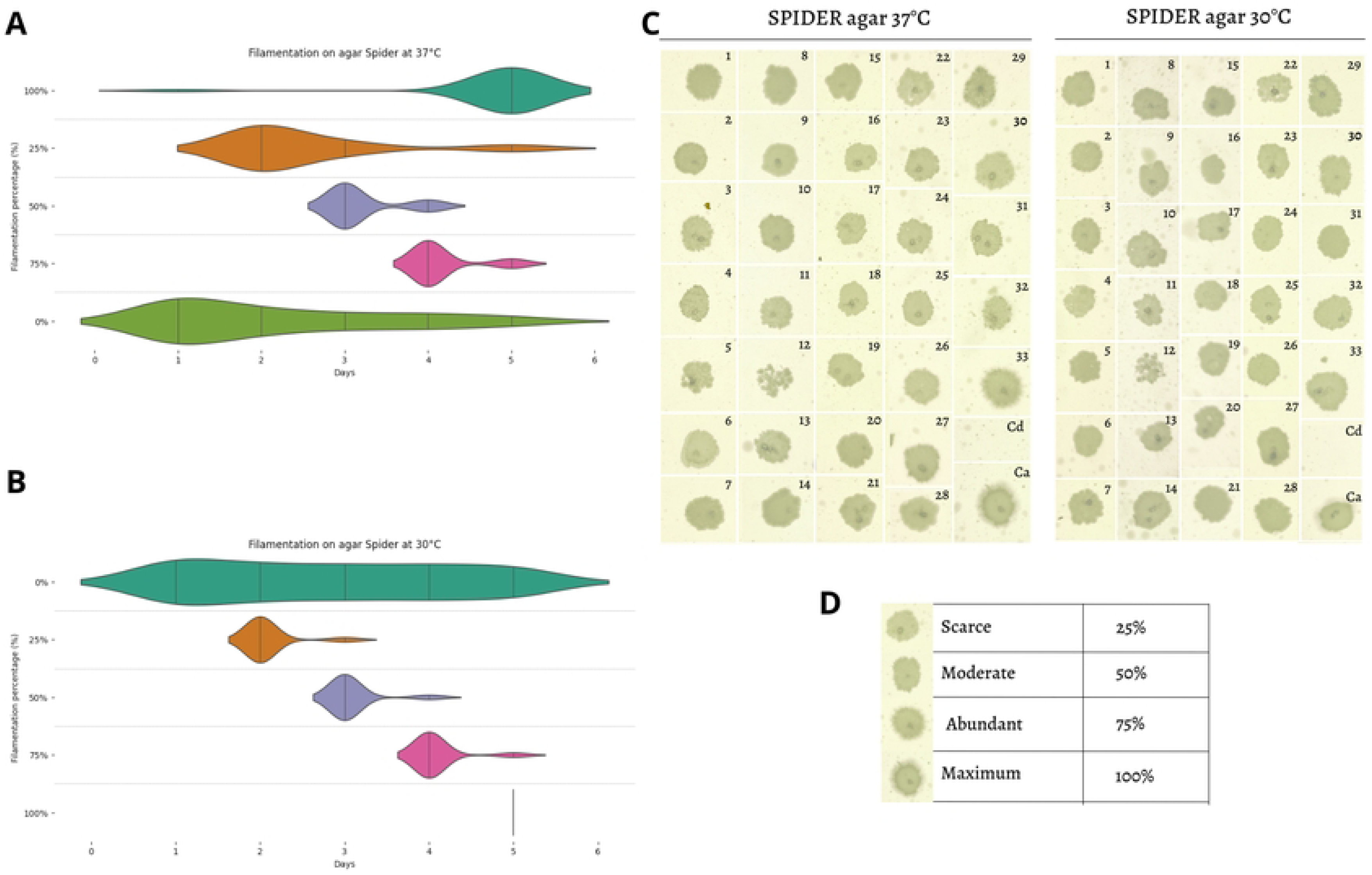
filamentation of growth in spots on Spider agar. A) Percentage of filamentation of isolates at 37°C based on the incubation period expressed in days. B) Percentage of filamentation of isolates at 30°C based on the incubation period expressed in days. C) Pictures of grown spots on spider agar at 37 and 30°C. Ca: *Candida albicans*, Cd: *Candida dubliniensis*. D) Filamentation capacity, percentages were assigned as follows: 25% scarce, 50% moderate, 75% abundant, and 100% maximum filamentation.

Finally, we tested antifungal susceptibility with the Sensititre™ YeastOne™ Y010 (SYO) panel, which follows criteria set forth by Clinical and Laboratory Standards Institute (CLSI) [42] and includes four azoles, three echinocandins, 5-Fluorocytosine, and Amphotericin B (**Table 1**). In the absence of established clinical breakpoints for *C. dubliniensis*, previously-defined epidemiological cutoff values (ECVs) were applied to identify isolates with decreased susceptibility that may harbor genetically encoded resistance mechanisms [43–45]. Moreover, four of the reduced susceptibility strains (Cd-4-6-16-22) also presented an atypical phenotype, characterized by limited growth under stress conditions in both liquid and solid media [22]. These observations suggest possible fitness trade-offs resulting from antifungal drug adaptation.

**Table 1:** Minimum inhibitory concentration (MIC) values for each isolation and antifungal according to SYO10 method. Values exceeding the epidemiological cutoff values (ECV) for echinocandins: anidulafungin, micafungin, caspofungin, are highlighted in yellow, for azoles: posaconazole, voriconazole, itraconazole, fluconazole are highlighted in pink and for Amphotericin B in blue. ND: No growth. The * indicates the patients who were exposed to antifungal treatment [22].

| Patients ID | Sample ID | Amphotericin B | Anidulafungin | Micafungin | Caspofungin | 5-Fluorocytosine | Posaconazole | Voriconazole | Itraconazole | Fluconazole |
| --- | --- | --- | --- | --- | --- | --- | --- | --- | --- | --- |
| 03-0-C | Cd-1 | 0.25 | 0.03 | 0.015 | 0.06 | ND | 0.015 | ND | 0.03 | 0.25 |
| 06-0-C | Cd-2 | 0.25 | 0.06 | 0.06 | **0.25** | ND | 0.008 | ND | 0.015 | 0.12 |
| 06-0-C | Cd-3 | **0.5** | 0.015 | 0.03 | 0.06 | ND | ND | ND | 0.12 | 0.25 |
| 06-0-C | Cd-4 | 0.25 | 0.03 | 0.015 | **0.12** | ND | **0.25** | **0.12** | **0.25** | **4** |
| 06-0-C | Cd-5 | 0.25 | 0.03 | 0.03 | **0.12** | ND | ND | ND | ND | 0.12 |
| 07-0-C* | Cd-6 | ND | **0.25** | **0.25** | **0.25** | 0.06 | **0.12** | **0.03** | 0.12 | **2** |
| 10-0-C* | Cd-7 | 0.25 | 0.06 | **0.12** | **0.5** | ND | 0.03 | **0.015** | 0.06 | **0.5** |
| 20-0-C* | Cd-8 | **0.5** | 0.03 | 0.015 | **0.12** | 0.06 | 0.008 | **0.015** | 0.015 | **0.5** |
| 20-0-C* | Cd-9 | **0.5** | 0.06 | 0.015 | **0.12** | 0.06 | 0.015 | 0.008 | 0.015 | **1** |
| 20-0-C* | Cd-10 | 0.25 | 0.06 | 0.06 | **0.25** | 0.06 | 0.03 | **0.03** | 0.06 | **2** |
| 22-0-C | Cd-11 | **0.5** | 0.06 | 0.03 | **0.12** | 0.25 | ND | ND | 0.015 | 0.25 |
| 22-0-C | Cd-12 | 0.25 | 0.06 | 0.06 | **0.12** | 0.06 | 0.015 | **0.015** | ND | **0.5** |
| 22-0-C | Cd-13 | 0.25 | 0.06 | 0.03 | **0.25** | 0.12 | 0.08 | ND | ND | 0.12 |
| 23-0-C | Cd-14 | 0.25 | 0.06 | 0.06 | **0.25** | 0.12 | 0.015 | ND | 0.015 | 0.25 |
| 23-0-C | Cd-15 | 0.25 | **1** | **0.12** | **0.25** | 0.25 | 0.03 | **0.06** | 0.03 | **1** |
| 24-0-C* | Cd-16 | 0.25 | 0.03 | 0.03 | **0.12** | ND | 0.008 | 0.008 | 0.015 | **1** |
| 24-0-C* | Cd-17 | 0.25 | 0.015 | 0.03 | 0.06 | ND | 0.015 | ND | 0.015 | 0.25 |
| 24-0-C* | Cd-18 | 0.25 | 0.015 | 0.015 | **0.12** | ND | 0.008 | ND | ND | 0.25 |
| 25-0-C* | Cd-19 | 0.25 | 0.015 | 0.03 | **0.12** | 0.015 | ND | ND | ND | 0.12 |
| 25-0-C* | Cd-20 | 0.25 | 0.06 | 0.03 | **0.12** | ND | 0.03 | **0.015** | 0.06 | **0.5** |
| 25-0-C* | Cd-21 | 0.25 | 0.06 | **0.12** | **2** | 0.25 | 0.06 | **0.015** | 0.06 | **1** |
| 25-0-C* | Cd-22 | ND | 0.06 | 0.06 | 0.06 | 0.06 | 0.06 | **0.03** | 0.06 | **2** |
| 25-0-C* | Cd-23 | 0.25 | 0.03 | **0.25** | **0.25** | ND | 0.008 | ND | ND | 0.25 |
| 25-0-C* | Cd-24 | **0.5** | 0.06 | 0.03 | **0.12** | 0.12 | 0.015 | ND | 0.03 | 0.25 |
| 25-0-C* | Cd-25 | 0.25 | 0.015 | 0.03 | **0.12** | ND | ND | ND | ND | 0.12 |
| 39-0-C* | Cd-26 | 0.25 | 0.03 | 0.03 | **0.12** | ND | 0.03 | **0.015** | 0.06 | 0.25 |
| 39-0-C* | Cd-27 | 0.25 | 0.06 | 0.03 | **0.12** | 0.06 | 0.008 | ND | 0.015 | 0.25 |
| 39-0-C* | Cd-28 | **0.5** | 0.06 | 0.03 | **0.12** | ND | 0.008 | ND | ND | 0.12 |
| 39-0-C* | Cd-29 | 0.25 | 0.06 | 0.03 | **0.12** | ND | 0.03 | **0.015** | 0.06 | 0.25 |
| 39-0-C* | Cd-30 | **0.5** | **0.12** | 0.06 | **0.25** | ND | 0.03 | **0.03** | 0.06 | **0.5** |
| 29-0-C | Cd-31 | **0.5** | **0.12** | 0.03 | **0.25** | ND | 0.03 | 0.008 | 0.12 | **0.5** |
| 50-0-C | Cd-32 | 0.25 | 0.03 | 0.015 | **0.12** | ND | 0.03 | 0.008 | 0.06 | **0.5** |
| 50-0-C | Cd-33 | **0.5** | 0.06 | 0.03 | **0.12** | ND | 0.03 | 0.015 | 0.06 | **0.5** |

Most of the analyzed isolates were below the CLSI-defined ECV for Amphotericin B, with nine (Cd-3-8-9-24-28-30-31-33) reaching that value. For 5-fluorocytosine (FC), no ECV has been established for this species. However, our data suggest high susceptibility, with 14 out of the 33 isolates having low MIC values and the remaining isolates being unable to grow under the test conditions. As for echinocandins, all but four isolates reached or surpassed the proposed ECVs in at least one echinocandin (**Table 1**). Specifically, isolates Cd-6 and Cd-15 were non-susceptible to all three echinocandins tested—Anidulafungin (AND), Micafungin (MF), and Caspofungin (CAS). In addition, isolates Cd-7, Cd-21 and Cd-23 were non-susceptible to MF and CAS, while isolates Cd-30 and Cd-31 were non-susceptible to AND and CAS. Although CLSI has not defined an ECV for CAS in *C. dubliniensis*, Espinel-Ingroff et al. (2015) [43] suggested a value of 0.12 μg/mL; under this criterion, only four strains were categorized as susceptible. For azoles, MIC values were often at or above proposed ECVs, especially for fluconazole (FZ) and voriconazole (VOR). Interestingly, resistance to these two azoles was significantly correlated, with 11 strains (out of 13 with low susceptibility to voriconazole, and 16 to fluconazole) showing low sensitivity to both drugs (P-value <0.005 in a Fisher’s exact test for the association). Itraconazole and posoconazole were generally effective, with only two and one strains with susceptibility at or below the ECV cutoff, respectively, and always in combination with low susceptibility to FZ and VOR.

### Genetic variability and novel genotypes revealed through MLST

To assess the genetic diversity within our collection of isolates, we first performed Multiple Locus Sequence Type (MLST) analysis using a previously proposed scheme comprising eight loci [44]. This identified the *MPI1* locus as the most informative, with nine different genotypes in the studied population, including one reported for the first time in this study (genotype 9 found in Cd-1). In contrast, *ACC1*, *ADP*1, *exVPS13*, and *exZWF1* loci presented no variability, with a single genotype for these genes detected in our collection. We combined information of all eight loci to assign a Diploid Sequence Type (DST) to each isolate, and compared them with previously reported DSTs from [44, 45], which encompass strains from multiple geographic regions. While McManus et al. (2008) reported 26 DSTs and Asadzadeh et al. (2017) added seven more (total 33 DSTs), our analysis identified six previously unreported DSTs, which were assigned numbers 34 to 39, following established nomenclature (**Supporting information Table ST1**). Notably, only one isolate in our collection belonged to an already described DST (Cd-1, DST 9), previously identified in France and Kuwait [44, 45]. One of the newly-reported DST (DST 34) was the most prevalent in our studied population (n=16, 51.6%), followed by DST 35 (n=8, 25.8%). These results expand the known genetic diversity of *C. dublinensis*, indicate a broad global distribution of DST 9, and point to a large genetic variability within the studied population, despite a narrow geographical origin. Phylogenetic analysis based on concatenated MLST loci from our collection grouped isolates into three main clades (**Fig. 4A**). Clade 2 predominantly contained DST 35 isolates, except for Cd-25, which clustered in Clade 1. Clade 3 mainly included DST 34 isolates, alongside other DST variants. Given the limited polymorphism at standard MLST loci, we expanded the analysis to the full-length sequences of the eight genes of the MLST scheme (*AAT1, ACC1, ADP1, MPI, RPN2, SYA1*/*ALA1, VPS13, ZWF1*) (**Fig. 4B**). Nucleotide changes resulting in amino acid substitutions were detected across all MLST loci except *ZWF1*. *SYA1* and *VPS13* showed the highest variability, with four amino acid substitutions each; *SYA1* exhibited a greater number of homozygous variants. (**Supporting information Table ST2**)

**Fig. 4:**
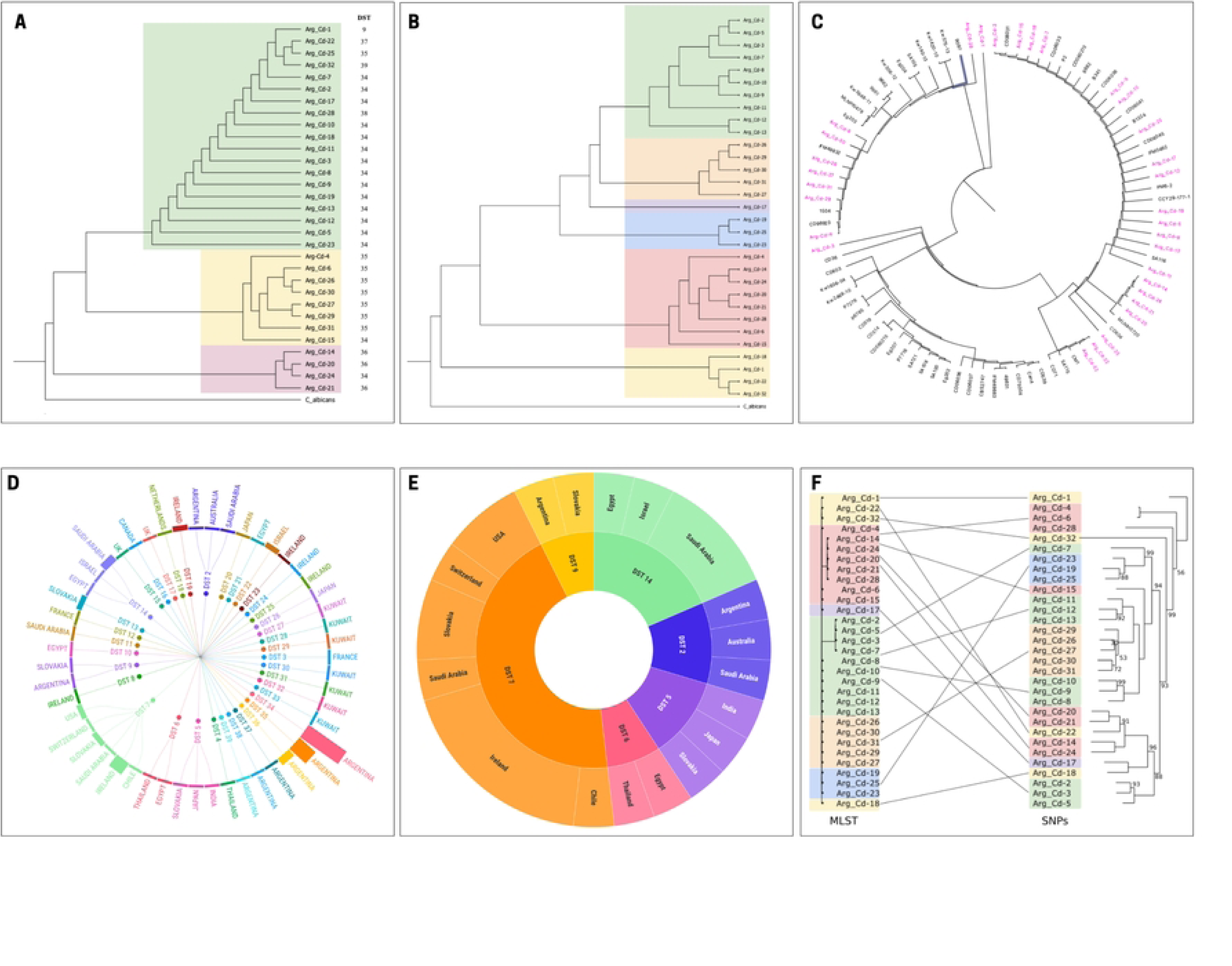
Maximum likelihood of phylogenetic trees based on MLST schemes. *Candida albicans* was used as an outgroup to root the tree. A) Based on the combination of the diploid sequence typing (DST) loci described by Mc Manus et al. 2018 [44]. Colors indicate clades: Clade 3 green, clade 2 yellow, and clade 1 pink. The clade 3 is the one that grouped most of the isolates. B) Based on the combined full sequences of the 8 MLST genes for *Candida dubliniensis*. Colors indicate clades: clade 1 yellow, clade 2 pink, clade 3 light blue, clade 4 violet, clade 5 orange, and clade 6 green. C) Combination of the DST loci from the 8 MLST genes in this study, indicated in pink, alongside *Candida dubliniensis* isolates from various geographical locations. Related to Supporting Information - Table ST1 D) Graphical representation of the DSTs grouped by color. Each node represents a DST, and each branch groups together those DSTs that are found in more than one country. E) Schematic representation of those DSTs that are found in more than one country and show similar phylogenetic relationships. F) Phylogenetic tree based on the current MLST schema on the left. On the right strain tree based on WGS data. Clades are coloured according to the clades found in the MLST based phylogenetic tree. Lines connect the same strain across the two trees. Support for the WGS is indicated when the bootstrap is below 100.

To explore population structure and geographic relationships, the concatenated sequences were compared with global isolates from previous studies [44, 45]. From the newly described DSTs of this study, DST 34 clustered close to DST 7, which has been previously found in diverse geographical locations such as Europe, the United States, Thailand, Chile, and Saudi Arabia. Certain isolates of DSTs 34, 35, and 36 showed closer affinity to strains from Thailand (DST 4) and France (DST 12). DST 37 and 39 isolates were phylogenetically linked to DST 2, while other DST 34 isolates related to strains from the Middle East, Japan, and Europe, such as the reference CD36 and strain CBS2747, indicating a potential global distribution. On the other hand, DST 35 is phylogenetically related to DST 5, which includes strains from Japan, Slovakia, and India. Likewise, DST 38 is related to different DSTs that belong to the Middle East and Slovakia. These results indicate that some DSTs are more localized in specific regions, while other genotypes are present in different geographical locations, suggesting a possible worldwide dispersion of these strains. (**Fig. 4C, 4D** and **4E, Supporting information Table ST3**)

### Genomic variability in *C. dubliniensis*

To characterize the genomic variability among the isolates, we sequenced their whole genomes with short reads and performed small variant analysis based on mapping the sequencing reads to the reference genome (see Materials and Methods). The number of detected SNPs between an isolate and the reference ranged between 11,370 and 17,942, suggesting a significant divergence to the reference strain (**Table 2**) All isolates exhibited variability in the distribution of genetic variants, and both homozygous and heterozygous sites were detected (average number of homozygous SNPs: 6680 with a range that goes from 4209 to 10812, average number of heterozygous SNPs: 9124 with a range that goes from 918 to 13361). Notably, isolates Cd-4 and Cd-6, characterized as atypical (see Table 5), displayed the lowest number of heterozygous variants (918 and 929 respectively). Two pairs of co-isolates that were presenting different colony size morphologies and were sequenced individually, Cd-1_SMALL and Cd-1_BIG, and Cd-5_SMOOTH and Cd-5_CREPE were genetically very similar, and were unified for the remainder as Cd-1 and Cd-2, respectively. (**Table 2, Supporting information Fig. SF16 and Table ST4**).

**Table 2:**
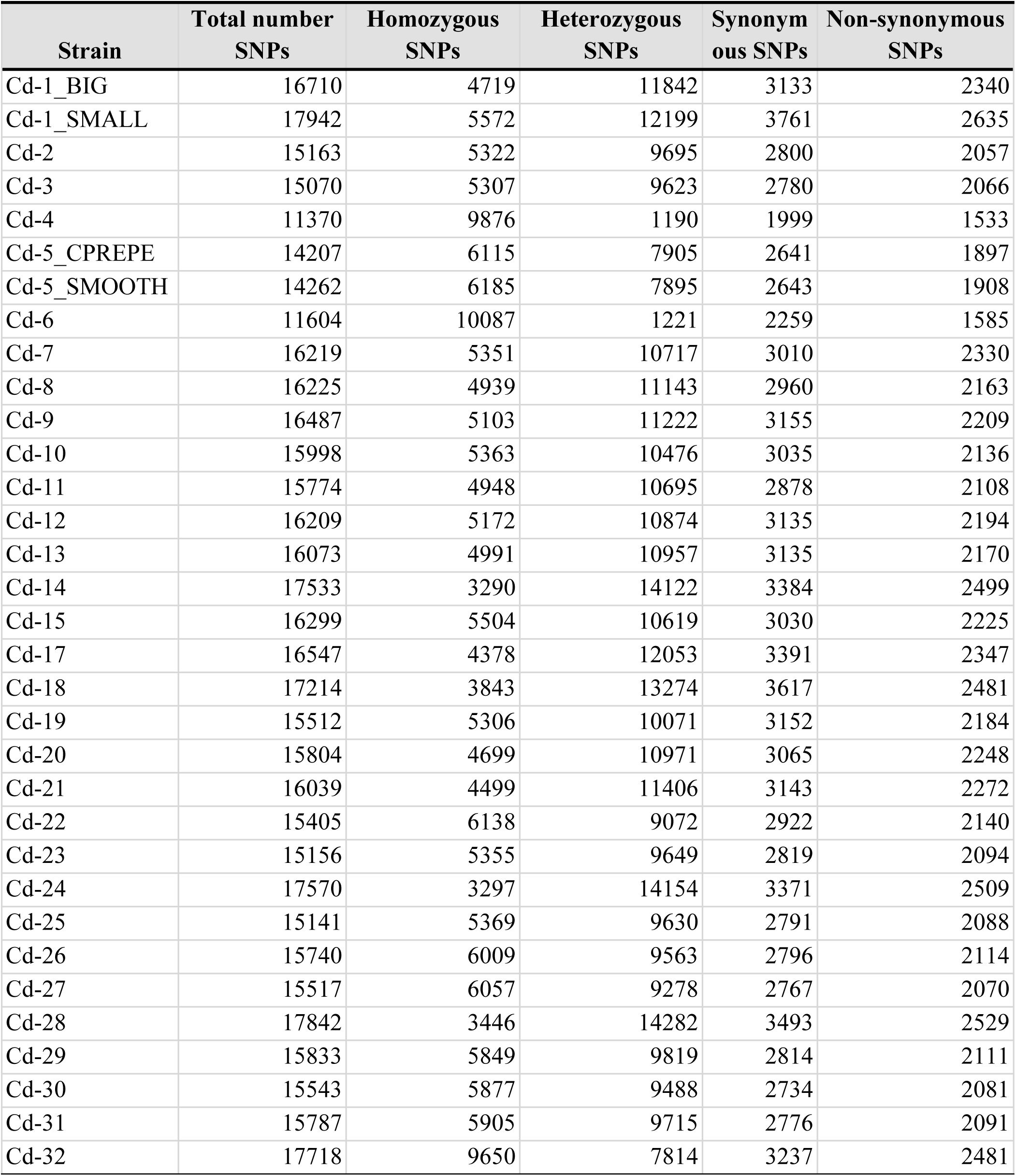
Number of detected SNPs between an isolate and the reference strain.

To further explore the genetic relationships among the isolates, a Multiple Correspondence Analysis (MCA) was conducted on their SNP profiles (**Supporting information Fig. SF17**). This analysis separated the isolates among two main axes collectively explaining 23.8% of the observed variability. The first axis separated strains depending on the number of homozygous and heterozygous SNPs with respect to the reference, separating Cd-32 (high number of both homozygous and heterozygous SNPs), and the two strains with low heterozygosity mentioned above (Cd-4 and Cd-6), from each other and the rest of strains. The second axis displays a gradient where several clusters of genetically-related (clonal) isolates can be observed. Notably, some of these clonal clusters comprised isolates from different patients. Conversely, we also identified that some isolates from the same patient, and even from the same periodontal site appeared distant from one another in the MCA, underscoring the potential for intra-host genetic diversity of resident *C. dublinensis* populations. (**Supporting information Fig. SF17**).

To investigate potential genetic determinants underlying the previously-described antifungal non-susceptible phenotypes, we focused on non-synonymous SNPs affecting genes associated with resistance (**Table 3, Supporting information Table ST5**). Homozygous and heterozygous non-synonymous variants in these genes were identified, enabling classification of strains as wild-type (WT) if no variants were present or non-WT if variants existed. Notably, a homozygous variant resulting in the loss of the stop codon in the *CDR1* gene (* /756Y) was detected in twelve isolates (Cd-4-6-15-20-21-22-26-27-29-30-31-32) of which most were resistant to one or more of the tested azoles (**Table 3. Supporting information Table ST5**).

**Table 3:**
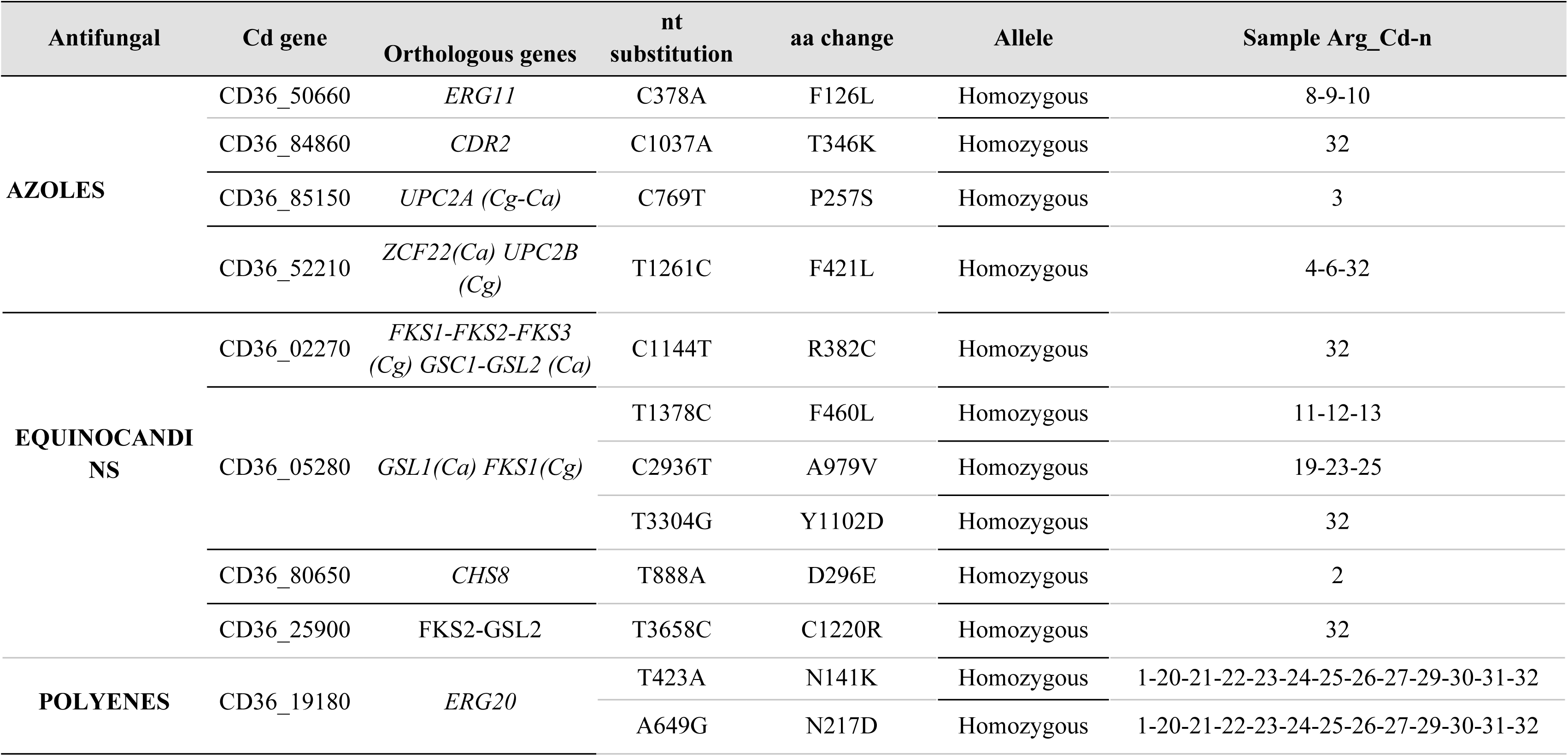
Homozygous non-synonymous variants related to antifungal resistance. Isolates are identified as Arg_Cd-n where “n” corresponds to each strain number. nt: nucleotide. aa: amino acid, Cd: *Candida dubliniensis*.

Three intra-patient isolates showing resistance to FZ (Cd-8-9-10) carried the F126L mutation in the *ERG11* gene, previously associated with FZ resistance in *C. albicans* [46]. Other isolates showing low susceptibility to some of the tested drugs carried homozygous or heterozygous mutations in genes previously associated with drug resistance (**Table 3, Supporting information Table ST5**). However, we could not find equivalent (homologous) mutations directly associated to drug resistance in the closest and well-studied relative *C. albicans* when searching for these mutations in the FungalAMR database [47]. Hence further research is required to establish causative links between these mutations and their potential role in the observed susceptibility phenotype.

### Development of a new genome-informed MLST scheme

Given the apparently low resolution provided by the existing MLST scheme (see above), we set out to assess the congruence of this scheme with genome-wide information from our collection, and evaluated the possibility of developing a more informative MLST. For this, we first reconstructed a genome-based phylogenetic tree of our isolated collection by building a pseudo-alignment in which the reference genome sequence was altered to represent the SNPs identified in each of the strains (see materials and methods). The resulting alignment included 25,631 variable positions, which were used to reconstruct a molecular phylogeny using a Maximum Likelihood approach (see materials and methods). This genome-wide phylogeny was compared to the MLST-based tree. This comparison (**Fig. 4F**) pointed to many inconsistencies between the two trees, and indicated that the genome-wide phylogeny better accounted for the diversity found in the considered *C. dubliniensis* strains. As seen in **Fig. 4F** the main clades found in the MLST tree were split in several different clades in the genome-wide tree. The genome-wide tree also displayed a more ramified structure, as compared to the MLST tree, indicating that the MLST scheme lacks resolution. In addition, some isolates that cluster together in the MLST tree are dispersed through very distant phylogenetic positions in the genome-wide tree, suggesting the MLST scheme inaccurately represents the evolutionary diversification of the strains. Considering these results, we explored the possibility of developing alternative MLST schemes that better represented the diversity found in *C. dubliniensis* as reflected in the genome-wide tree. To do so, we iteratively searched for sets of genes that, when concatenated and used to build a phylogeny, better reflected the genome-wide tree (see materials and methods). We obtained an optimal set of seven phylogenetic marker genes (**Table 4**) that were able to recover a tree with a normalized Robinson Foulds distance of 0.064 to the genome-wide tree, which corresponds to a single change in the tree topology. Adding additional genes to this MLST schema did not improve the tree, nor did any other combination of less than 10 genes. Hence, we propose that this alternative set of genes may provide a more accurate MLST scheme to chart the genetic diversity in *C. dublinensis*.

**Table 4:**
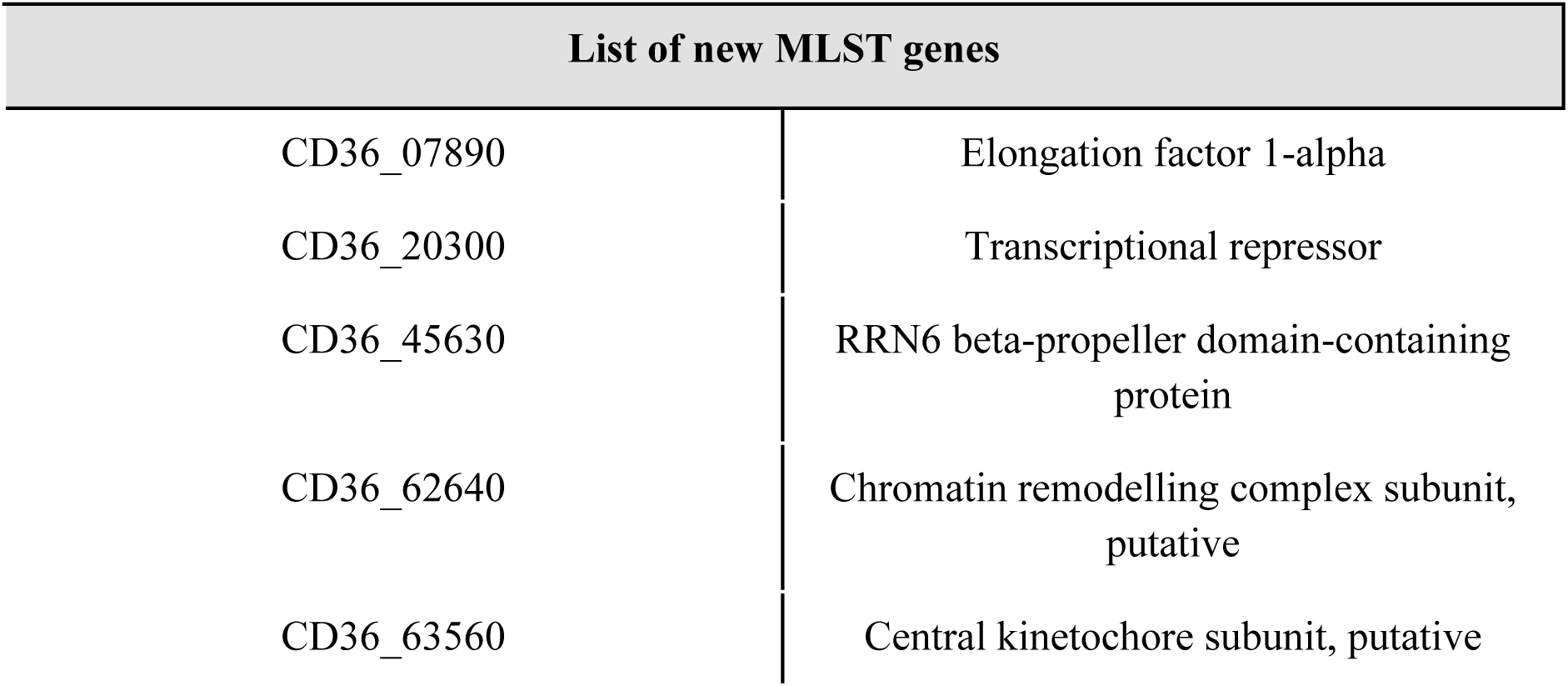
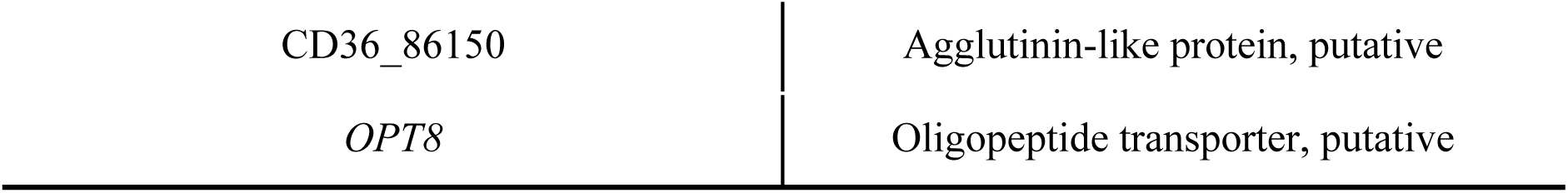
List of new MLST genes obtained to a set of MLST that better represented the diversity found in *Candida dubliniensis* as reflected in the phylogenetic tree reconstructed with the whole genome sequencing (WGS).

## DISCUSSION

In the last few years, C*. dubliniensis* has gained increasing attention from a clinical perspective. Although the incidence of this species in candidiasis remains comparatively low, the difficulty of distinguishing it from its closest relative *C. albicans* suggests that *C. dubliniensis* may have been underdiagnosed [22]. Similarly, as compared to *C. albicans,* our knowledge on the phenotypic and genetic diversity of worldwide isolates remains very limited. To address this gap, we extensively phenotyped and whole-genome sequenced, a collection of *C. dubliniensis* oral isolates from a well-defined clinical cohort of people living with HIV in Buenos Aires, Argentina [22]. To the best of our knowledge, this study represents the most extensive phenotypic and genomic characterization of *C. dubliniensis* isolates to date.

Unexpectedly, despite all isolates originating from the oral cavity and from a defined population in a specific restricted geographical origin, we observed a notable phenotypic diversity, even within pairs of samples co-isolated from the same patient and periodontal site. These findings suggest that *C. dubliniensis* displays notable phenotypic diversity and therefore varied potential to adapt to the oral niche, with marked differences in sensitivity to certain stresses. The magnitude of stress experienced by the fungus may vary depending on its commensal or pathogenic behavior. As a commensal, *C. dubliniensis* faces a mild immune response and a limited oxidative stress. However, in oropharyngeal candidiasis, a stronger immune response is mounted by the host, including neutrophil recruitment, which increases oxidative stress exposure [48]. We observed a large degree of variability in some morphological traits relevant for virulence such as the ability to filament, or the capacity to sustain oxidative stress. Consistent with previous research, our study supports that *C. dubliniensis* is generally more susceptible than *C. albicans* to high temperature, salinity, and oxidative stress [3, 48, 49–51]. These differences likely contribute to its reduced virulence, which is a multifactorial trait [51]. Similarly, we observed high osmotic stress sensitivity, which might be linked to the absence of sodium efflux pumps such as ENA21 [52] in *C. dubliniensis*. Growth under anaerobic conditions revealed that most strains grow and filament in this condition. When Spider medium was used, elevated temperatures enhanced filamentation. In the gingival sulcus and periodontal pocket, *C. dubliniensis* encounters hypoxic conditions, a significant environmental stressor [53]. In this regard, the periodontal pocket and crevicular fluid are considered to provide favorable environments for filamentation and growth in the hyphal form. [54].

We additionally investigated variability in susceptibility level to a panel of antifungal drugs commonly used to treat candidiasis. In the absence of clinical breakpoints we used previously proposed epidemiological cutoff values (ECVs), which serve as a sensitive marker of emerging resistance [55]. Several isolates exhibited MIC values above the proposed ECV for amphotericin-B, echinocandins and/or azoles. In contrast all isolates were highly susceptible to 5-Fluorocytosine. As for caspofungin, most isolates showed low susceptibility to this drug. However, as previously reported [43], susceptibility measurement for this drug is highly variable and unreliable. Sensitivity for the other echinocandins was generally below ECV levels, with a few cases clearly above these cutoffs. For azoles, low susceptibility was common, with more than 50% (18/33) of the strains having MIC values at or above ECVs for at least one of the tested azoles. Fluconazole was the azole with the largest number of strains with MIC at or above ECVs (16/33). Interestingly, low sensitivity to voriconazole and to fluconazole were often co-occurring. This may indicate common cross-resistance between the two drugs in *C. dubliniensis* [56]. Finally, 9 out of 33 strains showed MICs at the ECV cut-off for amphotericin-B. Despite its known nephrotoxicity, this drug remains a key treatment for invasive candidiasis in critically ill patients. Resistance to amphotericin-B in *Candida* spp. is rare [57] and is seldom observed in combination with resistance to other antifungals [58].

To understand the basis of the large phenotypic diversity observed in our collection, we first analyzed the genetic diversity by using the established MLST scheme for this species [44]. Our collection revealed previously unreported genotypes and DSTs, with only one shared genotype with previous studies. This aligns with prior research [44, 45, 59], indicating that the clinical population structure based only in MLST of *C. dubliniensis* is predominantly clonal, with limited microvariation. Such high clonality contrasts with the large phenotypic variation observed, suggesting the currently used MLST loci lack resolution. These findings suggested the need for a whole genome MLST strategy [60, 61], which may offer improved resolution and a more accurate representation of *C. dubliniensis* diversity, possibly unknown to date. We sequenced the whole genomes of all the strains in our collection, allowing us to propose a new MLST scheme, based on the exploration of the phylogenetic signal of the entire gene set of *C. dubliniensis*. The genome-wide tree revealed a more complex phylogenetic structure, indicating that the current MLST scheme fails to fully represent the genetic relationships among strains. By testing different gene combinations, we identified a set of loci that provided better resolution. These findings highlight the potential for a new MLST scheme based on WGS data to more effectively capture the genetic diversity of *C. dubliniensis*.

The availability of genome sequences allowed us to investigate the possible causes of low drug susceptibility identified in our collection. In some cases we could pin-point specific mutations that could explain the phenotype. For instance, we identified a F126L mutation in the *ERG11* gene, which was homozygous in three isolates (Cd-8-9-10) from the same patient previously treated with fluconazole for fungal diseases. Although this mutation has been previously described in *C. albicans* [62, 63] and *C. dubliniensis* [62], its direct association with an azole resistance phenotype has not yet been confirmed [62, 63]. Similarly, we found that isolates with MIC values at or above ECVs for amphotericin-B carried amino acid substitutions in ergosterol biosynthesis genes, including *ERG20, ERG24*, or *ERG6*. Further functional studies will be necessary to assess the biological relevance of these mutations and their actual contribution to antifungal resistance in this species.

### Conclusions

Our results provide evidence that *C. dubliniensis* encompasses significant genetic and phenotypic variability. This phenotypic diversity includes important differences in morphology, as well as stress and drug sensitivity phenotypes that can be relevant for the commensal or pathogenic life-style within different human niches, such as the oral cavity. Phenotypic and genetic divergence in our collection was surprisingly large when considering the restricted source of origin of the isolates, and included genetically divergent pairs of strains isolated from the same patient and oral site in the same day. This high intra-patient diversity might be common in persons living with HIV, as a result of multiple colonizations facilitated by a weakened immune system. Importantly, we demonstrate that the current MLST-based scheme provides very limited resolution to discriminate between strains that are genetically and phenotypically divergent, suggesting the worldwide and intra-patient diversity of *C. dubliniensis* isolates might have been under-estimated. We propose a genome-based MLST scheme, which together with the newly obtained genome sequences, will improve future population studies and the clinical monitoring of this relevant species.

## MATERIALS AND METHODS

### Study Population and Isolate Description

We analyzed a collection of 33 *C. dubliniensis* clinical isolates obtained from twelve persons living with Human Immunodeficiency Virus (HIV) who underwent gingival and periodontal parameter assessment to determine eubiosis, gingivitis or periodontitis status, and received antiretroviral therapy within the public healthcare system in Buenos Aires, Argentina. Among these donors, six had documented records for fungal opportunistic infections and antifungal treatments. Eight patients were diagnosed with periodontitis of varying stages and grades, and four had good gingival health. *C. dubliniensis* isolates were initially identified using phenotypic methods and subsequently confirmed by molecular techniques. In total, 36 isolates were confirmed as *C. dubliniensis*, of which 33 viable strains were selected for genomic analysis. Notably, four isolates displayed atypical colony morphologies (**Table 5**), which have been described elsewhere [22].

**Table 5:** Isolates of *Candida dubliniensis* used in this study. Those with atypical morphologies are marked with *.

| Sample ID | Patient ID | Site of isolation |
| --- | --- | --- |
| Arg_Cd-1 | 03-0-C | Subgingival biofilm |
| Arg_Cd-2 | 06-0-C | Subgingival biofilm |
| Arg_Cd-3 | 06-0-C | Subgingival biofilm |
| *Arg_Cd-4 | 06-0-C | Subgingival biofilm |
| Arg_Cd-5 | 06-0-C | Subgingival biofilm |
| *Arg_Cd-6 | 07-0-C | Subgingival biofilm |
| Arg_Cd-7 | 10-0-C | oral mucosa |
| Arg_Cd-8 | 20-0-C | Subgingival biofilm |
| Arg_Cd-9 | 20-0-C | Subgingival biofilm |
| Arg_Cd-10 | 20-0-C | Subgingival biofilm |
| Arg_Cd-11 | 22-0-C | Subgingival biofilm |
| Arg_Cd-12 | 22-0-C | Subgingival biofilm |
| Arg_Cd-13 | 22-0-C | Subgingival biofilm |
| Arg_Cd-14 | 23-0-C | Subgingival biofilm |
| Arg_Cd-15 | 23-0-C | oral mucosa |
| *Arg_Cd-16 | 24-0-C | Subgingival biofilm |
| Arg_Cd-17 | 24-0-C | Subgingival biofilm |
| Arg_Cd-18 | 24-0-C | Subgingival biofilm |
| Arg_Cd-19 | 25-0-C | Subgingival biofilm |
| Arg_Cd-20 | 25-0-C | oral mucosa |
| Arg_Cd-21 | 25-0-C | oral mucosa |
| *Arg_Cd-22 | 25-0-C | oral mucosa |
| Arg_Cd-23 | 25-0-C | Subgingival biofilm |
| Arg_Cd-24 | 25-0-C | Subgingival biofilm |
| Arg_Cd-25 | 25-0-C | Subgingival biofilm |
| Arg_Cd-26 | 39-0-C | Subgingival biofilm |
| Arg_Cd-27 | 39-0-C | Subgingival biofilm |
| Arg_Cd-28 | 39-0-C | oral mucosa |
| Arg_Cd-29 | 39-0-C | Subgingival biofilm |
| Arg_Cd-30 | 39-0-C | Subgingival biofilm |
| Arg_Cd-31 | 29-0-C | Subgingival biofilm |
| Arg_Cd-32 | 50-0-C | Subgingival biofilm |
| Arg_Cd-33 | 50-0-C | Subgingival biofilm |

### Phenotypic analyses and fitness assays in liquid medium

All 33 isolates were recovered from glycerol stocks, seeded on agar YPD (Yeast extract-Peptone-Dextrose), and grown for 48 h at 37°C. Single colonies were inoculated into 500 μL YPD broth in 96 deep-well plates (Nest Biotechnology, China) and incubated overnight at 37°C, agitating at 200 rpm with glass beads. Four technical and four biological replicates per isolate were performed. *Candida glabrata* CBS138, *C. albicans* CBS5314 and *C. dubliniensis* CBS7987 were used as positive controls; and uninoculated YPD medium as negative control.

After incubation, cell suspensions were diluted to 2-10 × 10^5^ (CFU/ml) in 96-well microplates (Nest Biotechnology, China). 3 μL of each dilution were inoculated into 197 μL YPD broth in 96-well microplates. Growth was recorded hourly over 24 h at 37°C, 200 rpm using a FilterMax F3 multimode microplate reader (Molecular Devices, USA) via optical density (OD) at 620 nm with SoftMax® Pro GxP Software (Molecular Devices, USA). The stress conditions tested included: oxidative (0.5 M H₂O₂) (Stanton, Argentina); redox (0.5 M Dithiothreitol (DTT)) (Bio-Rad Life Science, USA); cell wall stress (1 g/L Calcofluor-white (CFW)) and (30/25 µg/ml Congo Red (CR)) (SIGMA-ALDRICH, Germany); membrane stressor (100 µg/ml Sodium Dodecyl Sulfate (SDS)) (Biobasic Inc. USA); osmotic (0.75/0.5 M Sodium Chloride (NaCl) (Oxoid, UK), (1 M Sorbitol) and (6% Glycerol) (Anedra De Research AG, Argentina); protein denaturant (3% Absolute Ethanol) (Stanton, Germany) agents. [35, 64–69]. (**Fig. 5A**),

**Fig. 5:**
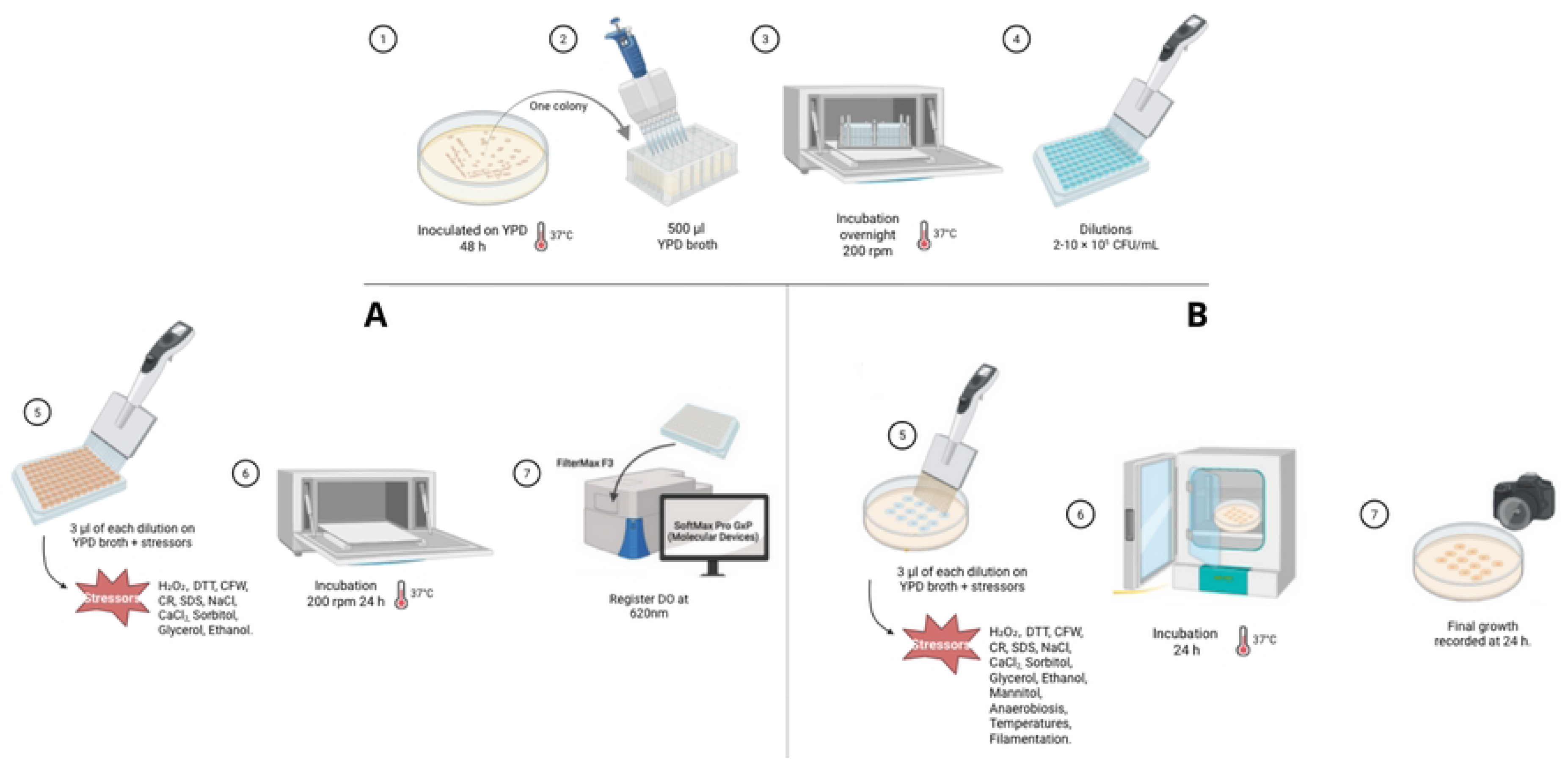
Schematic representation of the phenotypic fitness assays. 1. 33 *Candida dubliniensis* isolates, were seeded on agar YPD at 37°C for 2 days. 2. One single colony was inoculated into 500 μL of YPD broth, and 3. Incubated overnight with shaking (37°C, 200 rpm) and glass beads. 4. After incubation were diluted to 2-10 × 10^5^ (CFU/ml) before subsequent liquid (A) or solid (B) medium phenotypic assays were performed. **A)** Inoculation on broth YPD for growth curves: 5. After incubation, dilutions were performed. 3 μL of each dilution were inoculated into YPD broth. 6. Incubation was carried out at 37°C, 200 rpm, for 24 hours, with OD recorded every hour. 7. Growth was recorded by measuring optical density (OD) at a wavelength of 620 nm using a FilterMax F3 multimode microplate reader. **B)** Inoculation on agar YPD for spots test: 5. After incubation, dilutions were performed. 3 μL of each dilution were spot-inoculated onto Petri dishes. 6. Incubation was carried out at 37°C, for 24 hours. 7. Final growth on spots was photographed.

Data from OD measurements were used to construct growth curves, and statistical values were estimated using the Growthcurver R package (v0.2.1) [70]. This package fits a logistic growth model to the data and estimates key parameters including: k, n0, r, t_mid, t_gen, auc_l, auc_e, and sigma. For Principal Component Analysis (PCA), a subset of these parameters was used: k, n0, r, t_mid, t_gen, and auc_e, as they capture the most relevant aspects of growth dynamics under different conditions. Growth ratios were calculated for each stressor as the ratio: rAUC = AUC_strain_/AUC_YPD_ and relative to the WT strain *Candida dubliniensis* CBS7987: fAUC = AUC_stressor/_AUC_WT._

### Spot assays

Following the same growth and incubation conditions described above, stress response tests were conducted on YPD agar supplemented with chemical stressors at various concentrations. 3 μL of each dilution were spot-inoculated. The stress conditions tested included: oxidative (0.5 M H₂O₂); redox (0.5/1 M DTT); cell wall stress (1/2/5 g/L CFW) and (50/30/25 µg/ml CR); membrane stressor (100 µg/ml SDS); osmotic (0.75/0.5/1 M NaCl), (0.5M calcium chloride (CaCl_2_)) (Oxoid, UK) and (3/6% Glycerol); protein denaturant (3/5% Absolute Ethanol); (0.5/1 M Mannitol) and (1 M Sorbitol) without added glucose as cytoplasmic membrane stressors and as the unique carbon source. (1 M Sorbitol) with added glucose, solely as membrane stressors [35, 64–69]. (**Fig. 5B**)

Additionally, anaerobic growth was tested on YPD and Brucella blood agar with hemin (ASABRU) in an anaerobic jar (Oxoid, UK) with anaeropacks (Mitsubishi Gas Chemical, Japan), incubated for 7 days at 37°C [71]. Thermotolerance was assessed on YPD agar at 28, 30, 37, 42, 47°C for 5 days. Filamentation via mycelium formation was tested on Spider medium at 30 and 37°C for 5 days [35].

Final growth on spots was photographed and assigned numeric values from 0 to 4 to establish the observed growth level [72]: 0 = no growth, 1 = minimal growth, 2 = moderate growth, 3 = normal growth (similar to WT strain), 4 = overgrowth. Data were normalized by setting negative values to zero to eliminate inherent measurement bias, ensuring that negative values accurately represent the absence of growth.. According to the filamentation capacity, percentages were assigned as follows: 25% scarce, 50% moderate, 75% abundant, and 100% maximum filamentation.

### Statistical analysis

Statistical data processing was performed using R (v4.3.1) with packages “dplyr” (v1.1.3), [73], “ggplot2” (v3.4.3) [74], “reshape2” (v1.4.4) [75], “purrr” (v1.0.2) [76], “ggstatsplot” (v0.11.1) [77], and Python (v3.7.15) with “Pandas” (v1.3.5) [78], “Matplotlib” (v3.2.2) [79] and “Numpy” (v1.21.6) [80] in Google Colaboratory (Colab) [81].

### In vitro antifungal susceptibility

Antifungal susceptibility test was performed using the Sensititre™ YeastOne™ Y010 (SYO) colorimetric microdilution method (Thermo Fisher Scientific, USA), following the manufacturer’s protocol. Each microplate contains serial dilutions of ten dehydrated antifungals. From a 24-hour pure culture, 0.5 McFarland suspensions were prepared in sterile distilled water for each isolate. 20 μl of each suspension was inoculated into Sensititre culture medium to obtain a final concentration of 1.5–8 x 10^3^ CFU/ml. Subsequently, 100 μl of the inoculated broth were dispensed into each well of the SYO-microplates. To prevent contamination, each microplate was covered with a film provided by the manufacturer and incubated at 35°C for 24 h. Quality control strains *Candida parapsilosis* ATCC 22019 and *Candida Krusei* ATCC 6258 were included in each run following the Clinical and Laboratory Standards Institute (CLSI) guidelines [42]. Minimum inhibitory concentration (MIC) values were determined visually according to the color change and interpreted using the SYO panel reference and CLSI recommendations [42]. Given the lack of established Clinical Breakpoints (CBPs) for *C. dubliniensis*, epidemiological cutoff values (ECVs) were applied for interpretation, based on CLSI documents [43, 55, 82–84].

### Genomic DNA extraction and whole genome sequencing

Isolates were shipped to the Comparative Genomics Laboratory at the Institute for Research in Biomedicine (IRB), Barcelona. Strains were first grown on YPD (Yeast extract-Peptone-Dextrose) agar plates at 30°C for four days. We observed two different types of colonies (crepe versus smooth texture and small versus big colony size) in two out of 33 isolates. The other two isolates did not grow in these conditions. Cultures were grown from single selected colonies in an orbital shaker overnight (30°C, 200 rpm) in 15 ml of liquid YPD medium. Genomic DNA (gDNA) extraction was performed using the MasterPure Yeast DNA Purification Kit (Epicentre, France) following manufacturer’s instructions with slight modifications, and all reagents mentioned are from the kit if not specified otherwise. Briefly, cells were harvested using 3 ml of each culture by centrifugation at maximum speed for 2 min, lysed at 65°C for 15 min with 300 μl yeast cell lysis solution plus 1 μl RNAse A. After being on ice for 5 min, 150 μl of MPC protein precipitation reagent were added and centrifuged at 16.000 g for 10 min to pellet the cellular debris. The supernatant was transferred to a new tube and an extra RNAse treatment with RNAse A/T1 mix (Thermo Fisher Scientific, USA). Then a phenol-chloroform purification followed by a chloroform purification was done. Finally, DNA was precipitated using 100% cold ethanol, 0.1V of sodium acetate 3M and 2.5V of glycogen, and centrifuging the samples at 16.000 g, 30 min, 4°. The pellet was washed twice with 70% cold ethanol and, once the pellet was dried, the sample was resuspended in 100 μl of TE. Total DNA integrity and quantity of the samples were assessed by means of agarose gel, NanoDrop 1000 Spectrophotometer (Thermo Fisher Scientific, USA) and Qubit dsDNA BR assay kit (Thermo Fisher Scientific, USA).

Whole genome sequencing (WGS) was performed at the Centre Nacional d’Anàlisi Genòmica (CNAG). Libraries were prepared using the KAPA HyperPrep kit (Roche, USA) and quality checked on an Agilent 2100 Bioanalyzer (DNA 7500 assay). Sequencing was performed on an Illumina NovaSeq6000 in paired-end mode (150 bp reads). Read data was deposited in SRA under bioproject code PRJNA1347061.

### Genome analysis

Sequence reads from 31 isolates were processed to identify Single Nucleotide Polymorphisms (SNPs) using Freebayes [85], HaplotypeCaller [86], and Bcftools [87] as implemented in the PerSVade pipeline (v1.02) [88], using *C. dubliniensis* CD36 genome as reference (ASM2694v1). SNPs were filtered based on mapping quality (>30), QUAL value (>20), and read depth (>30). Only polymorphic sites supported by at least two callers were retained. A Multiple Correspondence Analysis (MCA) was conducted using ade4 R package dudi.acm function [89] and visualized with factoextra fviz_mca_ind function [90] to explore genetic relationships strains.

### Multilocus Sequence typing (MLST)

Eight housekeeping genes defined for *C. dubliniensis* MLST [44], were extracted by BLASTn alignment (v2.14.0) [91] against the reference genome *C. dubliniensis* CD36 ASM2694v1. The reference positions for each locus were extracted using bedtools (v2.30.0) [92] and sequences were reconstructed by replacing the reference nucleotide with the alternative SNP for heterozygous sites represented by the IUPAC ambiguity code. Sequences were assembled and aligned using MAFFT (v7.0) [93], concatenated and phylogenetic trees constructed with IQ-TREE (v1.6.12) under a GTR+G+I model with 1000 bootstraps [94]. Tree visualization was performed with iTOL (v6.8) [95]. Diploid sequence types (DSTs) were assigned following criteria from McManus (2008) and compared with published data. Since *C. dubliniensis* does not have a public MLST database, unlike other species in the genus, the same parameters established by McManus (2008) for the determination of diploid sequence types (DST) were followed [44]. Additionally, the generated DST sequences were compared with those previously published [44, 45].

### Comparative phylogenetic analysis

Pseudo-alignments were generated for each isolate by substituting SNPs into the reference genome, randomly selecting alleles at heterozygous sites. IQ-TREE (V2.1.2) [94] reconstructed 100 trees per alignment, which were concatenated to estimate bootstrap support. The trees were concatenated and the topology was compared to the one based on MLST. Bootstrap support was obtained by computing the number of reconstructed trees that contained a given clade.

### Development of MLST schemes

In order to find a set of genes that better represented the phylogenetic tree reconstructed using whole genome sequencing data (WGS), we substituted all SNPs on the reference genome, obtaining a pseudo-genome for each isolate. Positions containing heterozygous SNPs, indels or positions with a coverage below 20 were substituted by Ns in order to not consider them in the analysis while preserving their relative positions. Genes were extracted for each pseudo-genome and a pseudo multiple sequence alignment was built for each gene joining the sequences extracted from the different isolates. The multiple sequence alignment was used to reconstruct phylogenetic trees. To ensure enough variability was present, trees were only reconstructed for those multiple sequence alignments that contained at least six different sequences. This ensured that genes with little variability among strains were not considered. Phylogenetic trees for individual genes were then compared to the tree obtained based on the WGS. Both trees were rooted in the same leaf and then the normalized Robinson and Foulds (RF) distance was calculated using ETE v3 [96]. The 20 genes with the most similar trees based on the RF measure were selected and their alignment was concatenated to each of the single gene alignments. Trees were then built for each pair of concatenated genes and once again compared to the WGS tree. This process was repeated until ten genes were concatenated. The group with fewest genes giving the most similar tree to the WGS tree was the one kept.

